# Contact patterns reveal a stable dynamic community structure with fission-fusion dynamics in wild house mice

**DOI:** 10.1101/2020.02.24.963512

**Authors:** Jonas I. Liechti, Qian B., Barbara König, Sebastian Bonhoeffer

## Abstract

Living in groups is a widely adopted strategy in gregarious species. For group-living individuals it is crucial to be capable to integrate into a social structure. While there is an intuitive understanding that the concept of a group arises through some form of cohesion between its members, the exact definition of what constitutes a group and thus tasks like the detection of the dynamics of a group over time is a challenge. One way of measuring cohesion is through direct interactions between individuals. However, there is increasing evidence that associations between individuals can be mediated by others, and thus, that the drivers for group cohesion extend beyond direct individual interactions. We use dynamic community detection, allowing to relate individuals beyond direct contacts, both structurally and temporally, to study the social structure in a long-term study of a population of free-ranging house mice in a barn in Switzerland. During the 2-year study period, mice had unlimited access to food, and population density increased by 50%. Despite strong fluctuations in individual contact behaviour, population demography and structure embed into long-lived dynamic communities that are characterised by spatial fidelity, persist over several seasons and reproduction cycles, and considerably extend the life-span of single individuals. Within these multi-male and multi-female communities, seasonal changes strongly affect their structure, leading to fission-fusion like dynamics. We identify female-female interactions as the main driver for the longevity of these communities, a finding that contrasts with prior reports of the importance of a dominant male for the stability of a group. Moreover, males have a drastically shorter presence time in the study population and more often move between communities than females. Nevertheless, interacting with other breeding males in stable communities increases the duration of male presence and thus, potentially, reproductive success. Our analysis of contact patterns in a rodent that uses shelters to rest, hide and rear offspring emphasises the importance of female-bonded communities in the structuring of the population.

## 1 Introduction

Forming groups is one of the most widely spread adaptations of gregarious animals (Krause and Ruxton, 2002; Hayes and Schradin, 2017). Many species show variable social (as well as mating) systems, and only longer-term studies allowed to reveal common aspects to conclude on the social behaviour’s contribution to individual survival and reproductive success (for recent reviews see Clutton-Brock, 2016; Hayes and Schradin, 2017). However, detecting and describing adaptations of group living are not trivial tasks. Cohesion in groups extends beyond direct, pairwise relationships and depends on the ensemble of involved individuals (Sih *et al.*, 2009). Thus, relying on direct interactions between individuals as the basis for studies in social behaviour bares the risk of missing relevant aspects and may lead to an incomplete or distorted assessment about the factors relevant to living in groups. Traditionally, this dyadic approach formed the basis for studies of social behaviour in non-human animal systems. Scientists started only recently to consider indirect connections, predominantly by the use of social network theory (Wasserman and Faust, 1994), that allows to leverage network models as representations of the studied systems (Wey *et al.*, 2008; Croft *et al.*, 2011). Social network theory provided ample insights into the importance of indirect associations between individuals (for an overview see Brent, 2015; Krause *et al.*, 2015, 2009). A good example for the importance of indirect relations are the reported correlations between proximate fitness measures, such as individual survival or reproductive success, with centrality measures, like betweenness (Verdolin *et al.*, 2014; McDonald, 2007; Kanngiesser *et al.*, 2011; Gilby *et al.*, 2013) or eigenvector centrality (Stanton and Mann, 2012; Lusseau *et al.*, 2009; Lusseau and Newman, 2004).

A considerable part of social network analysis studies in non-human animals focuses on bigger mammals, e.g. dolphins (Lusseau, 2003; Stanton and Mann, 2012; Lusseau and Newman, 2004), elephants (Wittemyer *et al.*, 2005) or primates (Kanngiesser *et al.*, 2011; Ramos-Fernandez *et al.*, 2009; Flack *et al.*, 2006; Gilby *et al.*, 2013; Crofoot *et al.*, 2011). Specifically for small, crepuscular or nocturnal rodents, which played a key role in early behavioural and population structure studies (Bronson, 1979; Wolff and Sherman, 2008; Stenseth *et al.*, 1992), it is challenging to reveal principles of their social interactions by means of social network analysis (notable exceptions are Manno, 2008; Perkins *et al.*, 2009; Williamson *et al.*, 2016; Lopes *et al.*, 2016). House mice, *Mus musculus domesticus*, have been a recurrent study species in behavioural biology for decades, resulting in a rich literature on its social behaviour (see for example Bronson, 1979; Anderson and Hill, 1965; Crowcroft and Rowe, 1963; Singleton and Hay, 1983; Reimer and Petras, 1967; Wolff, 1985; Butler, 1980; Poole and Morgan, 1976). Males show territorial behaviour in the form of aggression towards other males (Singleton and Hay, 1983; Wolff, 1985). Albeit less pronounced, females have also been reported to contribute to territorial behaviour (Crowcroft and Rowe, 1963; Singleton and Hay, 1983; Butler, 1980), other authors find that females are less affected by territorial boundaries and thus able to move relatively freely between semi-natural enclosures (Crowcroft and Rowe, 1963; Baker, 1981; Manning *et al.*, 1995). Breeding can occur throughout the year under natural conditions, but is reduced during the cold months in winter (Bronson, 1979). Within groups, females are known to breed communally (König, 1994; Manning *et al.*, 1995) and it has been reported that aggression amongst females correlates inversely with the number of available mating partners (Rusu and Krackow, 2004), with both aspects being relevant only during breeding periods. The social system of the house mouse has therefore been summarized as being variable, with typically one dominant male monopolising reproduction with several females sharing his territory (Crowcroft and Rowe, 1963; Reimer and Petras, 1967; Bronson, 1979), suggesting that such social groups are demes. Others, however, reported much higher flexibility in females to move between territories (Lidicker, 1976), and increasing evidence based on genetic analyses does not support the concept of groups as demes. Baker (1981) reported common gene flow between “demes” and stated that the social organisation (social groups that defend territories) does not prevent gene flow within a population. Reporting of regular multiple paternity of litters further revealed that female house mice mate with males belonging to other groups (Baker, 1981; Stockley, 2003; Firman and Simmons, 2008; Ramm and Stockley, 2009; Manser *et al.*, 2011; Auclair *et al.*, 2014b; Thonhauser *et al.*, 2014). This illustrates pronounced variability in social behaviour in house mice. Long-term observations of populations are therefore required to learn about the principles that guide social interactions and allow to conclude on the benefits of variability in social interactions.

Most of what is known about the house mouse and its social organisation stems from direct visual observations, that lead to highly detailed descriptions of direct interactions between individuals. Limitations of this approach are the time consuming nature of visual observations and the absorption limit of the observer. Prolonged tracking of a population cannot easily be carried out both systematically and continuously. Nevertheless, already through prolonged visual observations, social groups have been found to be relatively stable, lasting over 100 days (Singleton and Hay, 1983; Reimer and Petras, 1967). Reimer and Petras (1967) reported evidence that within these stable groups the dominant male might be replaced by its offspring, leaving the group intact and suggesting a dominant role of males for the stability of a group. With the rapid progression of technical equipment and the development of methods that permit to process and analyse automatically generated data, studies of population structures are more often realised by means of automated tracking of individuals. Automated observations are advantageous in that they are less labor intensive and applicable in a systematic manner, besides being little invasive for the study animal. The disadvantage of automation manifests in the simplicity of single observations that often consist solely of physical proximity information. With automated tracking becoming more and more abundant, also temporal network analysis (Pinter-Wollman *et al.*, 2013; Holme and Saramäki, 2012; Holme, 2015; Dakiche *et al.*, 2019), an extension of network analysis that aims to efficiently include the temporal resolution of data, has made its way into the study of non-human animal societies.

Here we present a new approach to analyse the dynamics of the social structure of a free-ranging population of house mice over several seasons and years, with the aim to reveal principle aspects in individual behaviour that underlie the interaction pattern observed. To do so, we took advantage of the fact that house mice use rather small nesting places to rest and hide. Sharing such nest sites, therefore, allows to draw conclusions on the kind of social relationship among individuals (we do not expect individuals to spend more than a few seconds in the same nest if they interact agonistically) and on group membership (see Auclair *et al.*, 2014a; König and Lindholm, 2012; Lopes *et al.*, 2016). We carried out a social network based study of the structural dynamics within a population of house mice inhabiting a barn near Zurich, Switzerland. The set-up mimics natural conditions of house mice in middle Europe, providing shelter, food and the possibility for migration. As basis for our study we use automated observations in the form of directional movement information drawn from RFID-antennas (König *et al.*, 2015) at the entrance of artificial nest boxes used by the mice for resting and as safe places to rear litters. In a first part, we assess the composition of the studied population and characterize dyadic contact patterns. In a second part, we use dyadic contacts to construct a sequence of contact networks and describe the population structure under this network paradigm by means of social contact groups. This approach takes into account indirect associations between individuals for the determination of social groups. In a third part, we focus on the temporal dynamic of these social groups leveraging on dynamic community detection, i.e. we construct dynamic communities by following social groups through time, and ascertain what type of dyadic contacts determine the observed temporal patterns. Finally, we correlate properties of these dynamic communities with characteristics of associated individuals.

Our findings from the first and second part indicate the presence of strong fluctuations in the population structure, its demography, as well as in the individual contact behaviour. However, these fluctuations integrate into dynamic communities, described in the third part, leading to a remarkably stable social structure with strong spatial fidelity.

## 2 Results

### 2.1 Demography of the house mouse population

The studied population of wild house mice inhabits a barn near Zürich, Switzerland. Within the barn, an RFID-tracking system is set up, monitoring 40 artificial nest boxes. Fig. 1c depicts a schematic representation of a single nest box and its antenna set-up. The inside of the barn is depicted in Fig. 1a and a schematic overview with the approximative locations of the nest boxes is provided in Fig. 1b. All individuals of at least 18 grams are tagged (by implantation of a transponder), rendering them detectable by the tracking system (see Section 4.1 for details). In the following we will consider the collected data in a binned form as a sequence of time-windows of 14 days. We use the term *slice* to refer to a single aggregation window within this sequence (see Section 4.2 for details). The RFID-tracking system registered 448 individuals from January 2008 to January 2010. 346 of them were present for at least one month, i.e. more than two aggregation cycles. While we can assess the total presence time of individuals as the time span during which they were detected by the RFID-tracking system, the collected data does not allow to directly deduce life-span. As such, we hereafter only refer to the time of presence. An individual is on average present for 156.1 (standard deviation, *sd* = 133.8) days, with an average presence time of 8.5 (*sd* = 7.5) aggregation periods. This is a lower estimate because the presented data is gathered over two years only, while the house mouse population extends its existence over a far bigger time range, leading to censoring, a fact that will be taken into account in the survival analysis in Section 2.6.

**Fig. 1.**
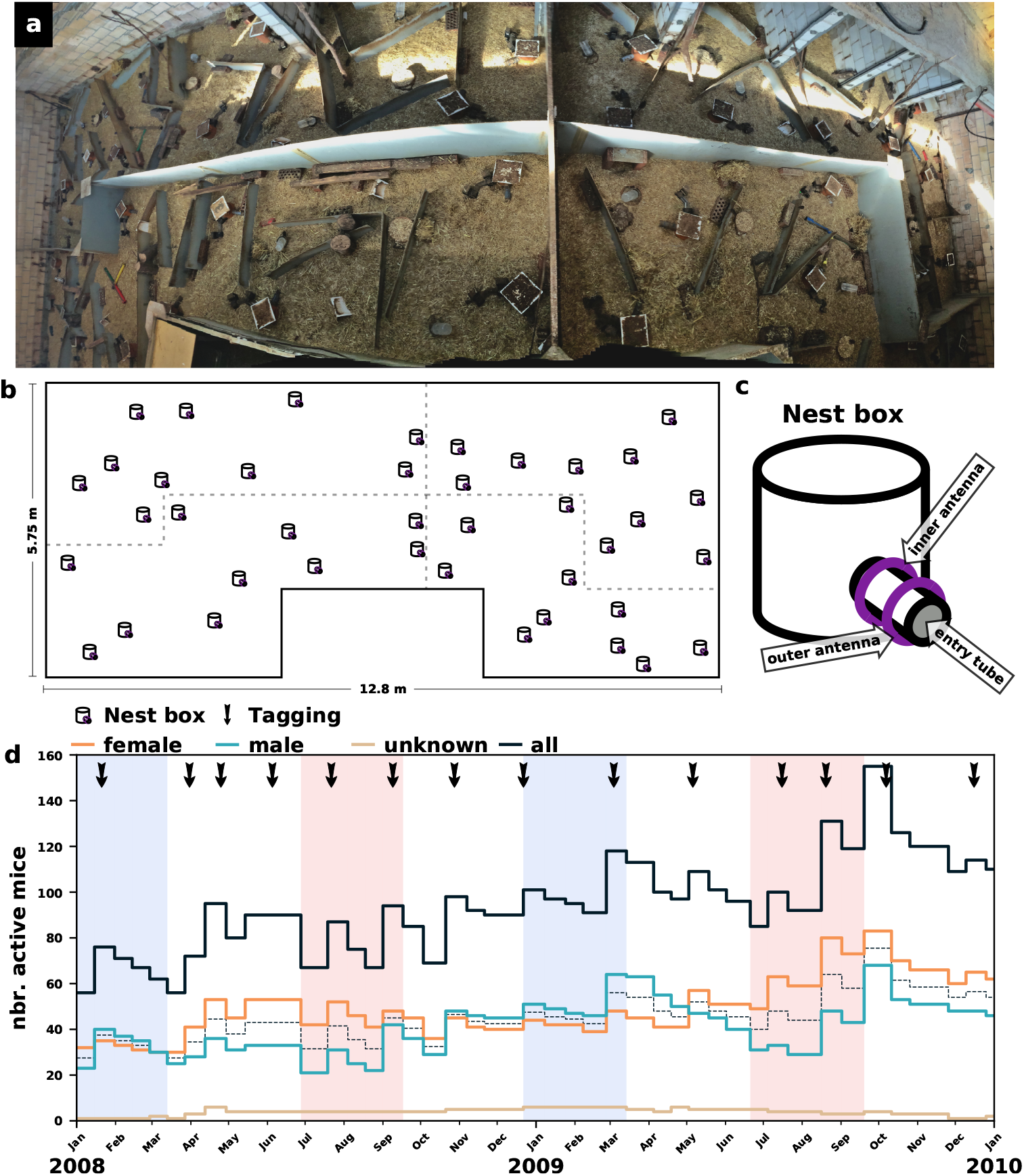
Illustration of the study system, a population of *Mus musculus domesticus* in a barn near Zürich, CH. **a**, Picture of the inside of the barn housing the population. Small, traversable walls (in gray) separate the foraging area into 4 zones which mice can access through several holes. Artificially placed obstacles further increase the complexity of the landscape in each zone. **b**, Schematic representation of the barn. Dotted lines indicate the separation of the foraging area into 4 zones. Black symbols indicate the approximate locations of the 40 artificial nest boxes. **c**, Schematic representation of a nest box. The entry tube provides exclusive access to the inside of the nest box and is equipped with an inner and an outer antenna (purple rings) allowing to detect directional movements (for details see König *et al.*, 2015). **d**, Time-series of the number of house mice detected by the RFID-readers during consecutive observation windows of 2 weeks. Single observation windows can be extended by periods of no data collection in order to assure an equal duration of data collection for each observation. See FIG. S1a for a version of this panel where periods of no data collection are highlighted. Blue and red areas designate winter and summer periods. Black arrows indicate tagging events (for explanation see Section 4). The dashed line indicates 50% of present individuals of known sex.

The total number of individuals over time, additionally separated for females and males, is shown in Fig. 1d. In summer, there are more females, while in winter there are more males (refer to Section 4.3 for a definition of summer/winter). The average sex-ratio (male/female) for the summer periods is with 0.59 (*sd* = 0.10) roughly half than that in winter, 1.11 (*sd* = 0.15), which corresponds to a significant shift of the sex ratio (*Kolmogorov-Smirnov*, *D*(10, 10) = 0.9, *p* < 0.01). The winter and summer periods considered are illustrated in red and blue in Fig. 1d (see Section 4 for details).

### 2.2 Social behaviour measured via dyadic contacts

Fig. 2a illustrates how we deduce pairwise contacts from the data provided by the RFID-tracking system. In Fig. 2b we report the average number of hours a pair of individuals from different categories (female, male, all) are meeting in nest boxes during an aggregation period. Similar to the pattern observed for the sex-ratio, the average contact intensity reveals a pronounced seasonal dependency transcending all categories. With a roughly nine-fold increase in average contact time from summer to winter, the male-male contacts are most affected by seasonality. This is followed by an approximately seven-fold change for female-male contact time and an almost four-fold increase in female-female contact time. This change is significant for all categories (*t-test*, *df* = 18 *throughout, all: t* = 5.1, *p* < 0.01*; female-female: t* = 6.3, *p* < 0.01*; male-male: t* = 4.9, *p* < 0.01*; female-male: t* = 4.9, *p* < 0.01). Note, that the category *all* is the only category to include contact pairs containing individuals with unknown sex.

**Fig. 2.**
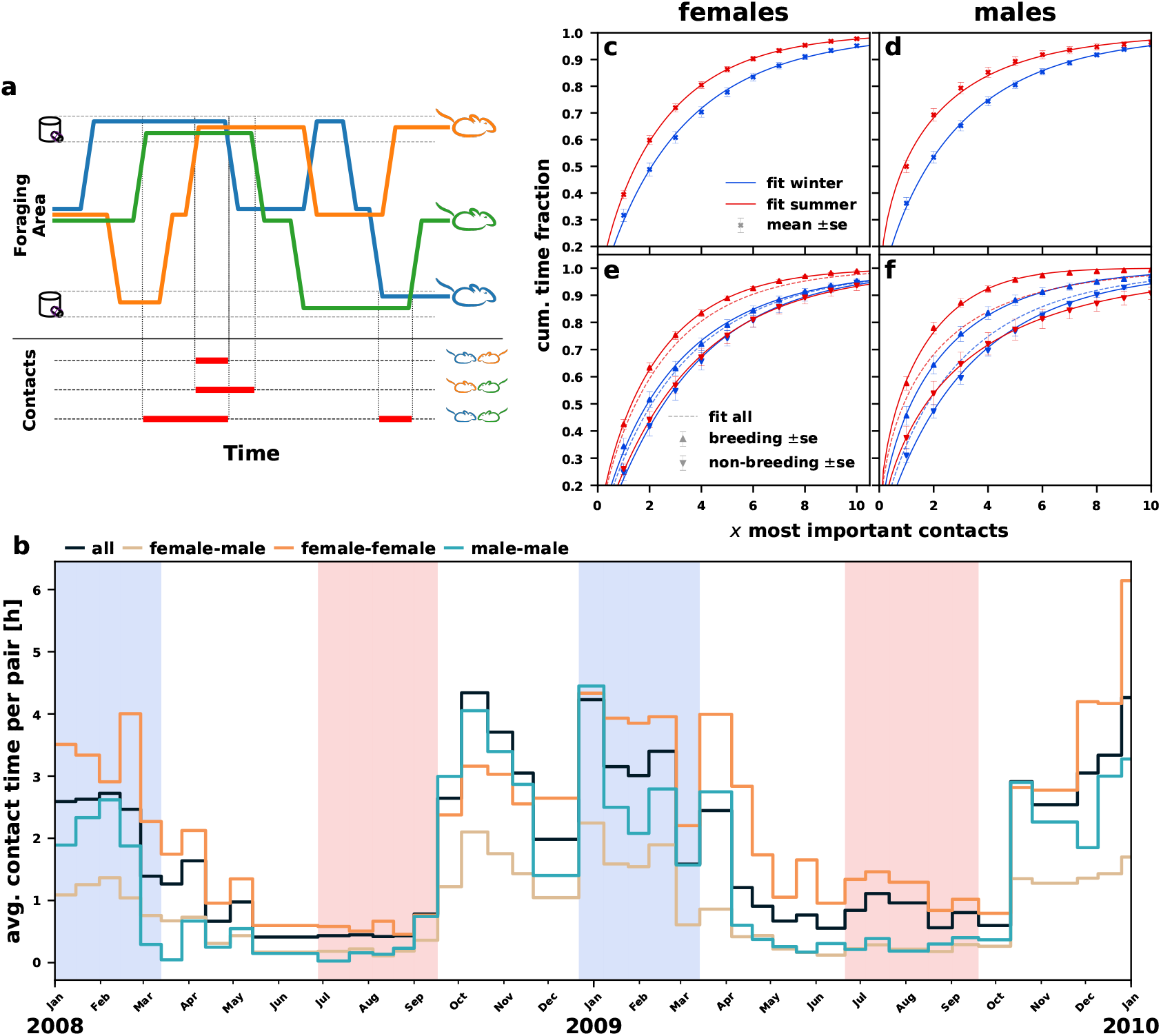
Pairwise contact data in a population of *Mus musculus domesticus*. **a**, Illustration of how concurrent visits of nest boxes lead to contacts between individuals. The time two individuals spend together in a nest box is considered as contact time. Concurrent stays create transitive contacts, i.e. the presence of two or more individuals leads to pairwise contacts for all combinations of pairs. **b**, Average contact duration between individuals from different categories. Blue and red areas designate winter and summer periods. See FIG. S1b for a version of this panel where periods of no data collection are highlighted. **c** to **f**, Cumulative distributions of the fraction of contact time assigned to contact partners, ranked from most important to least important, in summer (red) and winter (blue). Each measure (marker) designates an average, with the corresponding standard error as error-bars. The lines correspond to a fit of the raw data using an exponentiated Weibull distribution. **c** and **d**, Distribution of contact times of females and males. **e** and **f**, Distribution of contact time for breeding (up-warded triangles) and non-breeding (down-warded triangles) individuals. The dashed lines correspond to the fit from panels (c) and (d), respectively.

In Fig. 2c-f, we show the cumulative contact time-fraction attributed to contact partners, ranked from most to least important (see Section 4.3 for details). We report for an average female and male, both in summer and winter, including the breeding status in Fig. 2e and f. Note, that all cumulative distributions are well fitted by an exponentiated Weibull curve. To assess how time allocation changes between seasons, we test up to which rank in the distribution of contact time-fraction there is a significant difference between summer and winter. We observe a general trend of increased specificity during summer, i.e. a fixed fraction of contact time is distributed over fewer contact partners (see Fig. 2c and Fig. 2d). For males, this difference is significant up to the 9th rank (*t-test*, *df* = 213, *p* = 0.16) and for females up to the 13th rank (*t-test*, *df* = 228, *p* = 0.15). Including the breeding status yields an interesting insight: for both breeding males and breeding females, the seasonal changes seem more pronounced, as compared to the non-breeding individuals (see Fig. 2e and Fig. 2f). In fact, there is no statistical evidence that non-breeding individuals allocate their contact differently between seasons. This holds true both for males (1st rank: *t-test*, *df* = 112, *p* = 0.17) and females (1st rank: *t-test*, *df* = 64, *p* = 0.73). We conclude that between seasons only breeding individuals change contact behaviour significantly, such that they allocate contact time more specifically during the summer.

### 2.3 Population structure and social group dynamics

Based on all dyadic contacts we construct a sequence of social contact networks. We assess the group structure of the population at each slice in this sequence using group detection within the corresponding contact network. Based on the resulting sequence of group structures we deduce dynamic communities by tracking groups (or sets of groups) over multiple slices within this sequence. The dynamic community detection is carried out by the method by Liechti and Bonhoeffer (2019). This method is unique in the sense that it can detect dynamic communities with internal fission-fusion dynamics. Further details about how static networks are generated, how the social group structure is assessed and the dynamic communities are identified can be found in Section 4.

Considering the social group structure we find that, on average, the networks consist of 13.2 (*sd* = 2.9) groups in summer and only 8.9 (*sd* = 2.3) in winter which is a significant decrease (*Mann-Whitney*, *U* = 90, *p* < 0.01). Visualisations of the group structures in four randomly selected networks, one per season and year can be found in Fig. S4. We use the modularity score (Newman and Girvan, 2004), a standard measure in network analysis, to obtain a quantitative measure of how strong the networks are partitioned into social groups. The modularity of a group structure increases if more connections occur between individuals belonging to the same group. Thus, comparing the modularity score between two different group structures, or the group and the community structure, allows to deduce which structure better contains contacts within groups or dynamic communities. We find no significant evidence that winter periods score differently in terms of the modularity score when compared to group structures in summer (*Mann-Whitney*, *U* = 70, *p* = 0.14). Thus, while observing more groups in summer, we cannot conclude that the group structure would also be more pronounced. Spatially, the group structure depicts a high consistency in the sense that groups also tend to aggregate in space (see Fig. S3f-i).

Fig. 3a-d show visualisations of the dynamic community structure in four randomly selected networks, one per season and year (the same as in Fig. S4). The node positions are given by a ‘spring’-layout (Fruchterman and Reingold, 1991) which aims to decrease the distance between strongly connected nodes. The contours delineating the different dynamic communities are in good agreement with the positioning of the nodes, indicating that strongly connected nodes tend to belong to the same community. A more extensive version of this figure is available in Fig. S3 including the group structure through colouring of the nodes.

**Fig. 3.**
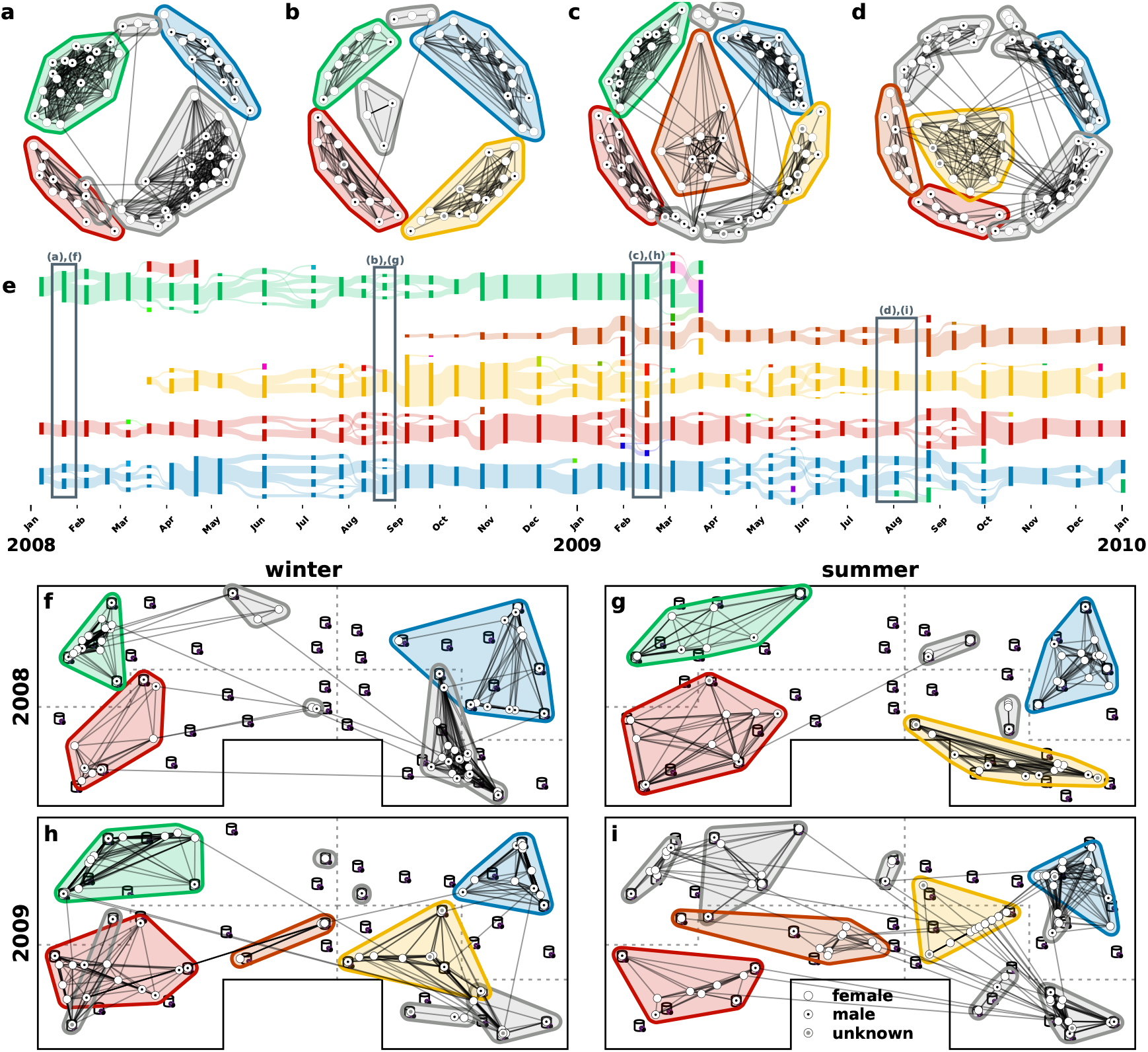
Temporal and spatial representations of the contact structure. **a** to **d**, Representations of the contact network at randomly chosen time points during the summer and the winter periods. The layout is deduced using the force-directed spring model by Fruchterman and Reingold (1991), implemented by Hagberg *et al.* (2008), that aims to position strongly connected nodes close to each other. The contours represent the dynamic community structure. The colouring of the contours is consistent throughout the figure with the life-cycle of the coloured dynamic communities depicted in panel (e). Grayed out are dynamic communities that do not figure in panel (e). **e**, Alluvial representation of the five most long-lived dynamic communities. Each rectangular block represent a group at a given time point with the colour indicating the dynamic community. The fluxes between groups visualize how the individuals regroup from one time point to the next. We can observe how dynamic communities fission into several sub-groups during summer that recombine during the cold winter period. The social and spatial illustrations depicted in the other panels are marked with a rectangular box. **f** to **i**, Illustration of contact structures embedded within the barn. Individual house mice, i.e. dots, are placed at their average position in the barn during the aggregation period, i.e. barycentre of the nest boxes visited weighted by the time spent in each box. For a more detailed version of this figure refer to FIG. S3. A version illustrating the group structure can be found in FIG. S4.

Fig. 3f-i illustrate the same networks but place each mouse at its average position in the barn. Since we can localize mice only via nest box visits, the average position is given by the weighted barycentre of the nest boxes they frequented, using the total time spent in nest boxes as weights. Localising mice this way, and thus embedding the social network into the barn, provides insights on how individuals distribute over the nest boxes and how social contacts relate to spatial proximity. We observe a clear pattern that social contacts predominantly occur between spatially closely located individuals. An animation illustrating the displacements of tagged mice, as well as the contacts between them, over the entire observation period is available in the supplementary materials (see *barn anim.mp4*).

### 2.4 Persistence in dynamic community structure

Fig. 3e illustrates the life span of the five most long lived dynamic communities within this population (for a representation of the entire dynamic community structure refer to Fig. S3e). We find two dynamic communities that exist over the entire observation period. This is particularly striking when considering that none of the initial members of these two dynamic communities are still present at the end of the observation period.

Quantitatively, the average life-span of the detected dynamic communities is 7.1 (*sd* = 10.3) slices. Here, we need to consider that the arithmetic mean can give a wrong impression: Over a fixed duration one might observe many more short-lived dynamic communities simply because more short-lived dynamic communities fit into the observation period. Therefore, we also report the life-span of the average community encountered by an individual. This weighted average is defined by the fraction of presence time an individual spent in a dynamic community of given life-span, summed over all individuals. The weighted average life-span of a community is with 29.0 slices considerably longer than the arithmetic mean. Even more importantly, it extends the average presence time of an individual by roughly 2.5 times.

As opposed to the group structure, we get no statistical evidence for seasonal patterns in the average number of dynamic communities (*Mann-Whitney*, *U* = 62.5, *p* = 0.36). This is in agreement with the observation that a dynamic community consists on average of 1.1 (*sd* = 0.3) groups in winter and 1.5 (*sd* = 0.7) groups in summer. Considering that the weighted average life-span of a dynamic community extends a year, we conclude that these seasonal changes in the group structure happen predominantly within dynamic communities. As such, they present community-internal fission-fusion dynamics. This internal decomposition of dynamic communities into sub-groups during the summer periods can also be observed visually in the life-cycles of the five dynamic communities depicted in Fig. 3e.

When comparing the positioning of dynamic communities between time points (contours in Fig. 3f-i), we conclude that none of the dynamic communities present in several panels drastically changed their location in the barn. Most remarkably, this holds true for the dynamic communities that persist throughout the 2 years observation period. Note, that the boxes occupied by a group during an aggregation window, as well as by dynamic communities over time, predominantly belong to a single one of the four areas. However, visual observations inside the barn give no indication that the borders between areas would represent an obstacle that is physically difficult for a house mouse to cross.

### 2.5 Composition of the dynamic community structure

To assess what type of dyadic contacts are most important for the formation of these long-lived dynamic communities, we consider the life-spans of dynamic communities in contact networks with parts of the contacts omitted. In particular, we consider the series of contact networks, once ignoring, and once exclusively considering same sex contacts (see Fig. S2 for a visualisation of the resulting dynamic community structures). Fig. 4a shows non-parametric fits (Kaplan and Meier, 1958) for the survival data of the resulting dynamic community structures. The male-male only contact structure leads to a significantly different distribution of life-spans (see caption of Fig. 4a for details), as compared to the total contact structure. Interestingly, female-female contacts produce dynamic communities with life-spans comparable to the ones of the entire population. When considering inter-sex connections, the trend is similar, although less pronounced. Ignoring male-male connections leads to long-lived structures that are comparable in terms of life-spans to the ones observed within the total population. If female-female connections are ignored, there is a non-significant trend that the resulting structures have a different pattern in terms of life-spans (*p=0.06*). These observations indicate that female-female contacts play an important role in the longevity of the observed dynamic communities within the population. Female-female associations thus are crucial for the stability of dynamic communities within the population. Our observations show that females generally do not move much. In fact, we find that during our observations only 23% of the females ever belonged to more than one community, whereas for males this fraction is, with 36%, significantly higher (*z* = −3.0, *p* < 0.01).

**Fig. 4.**
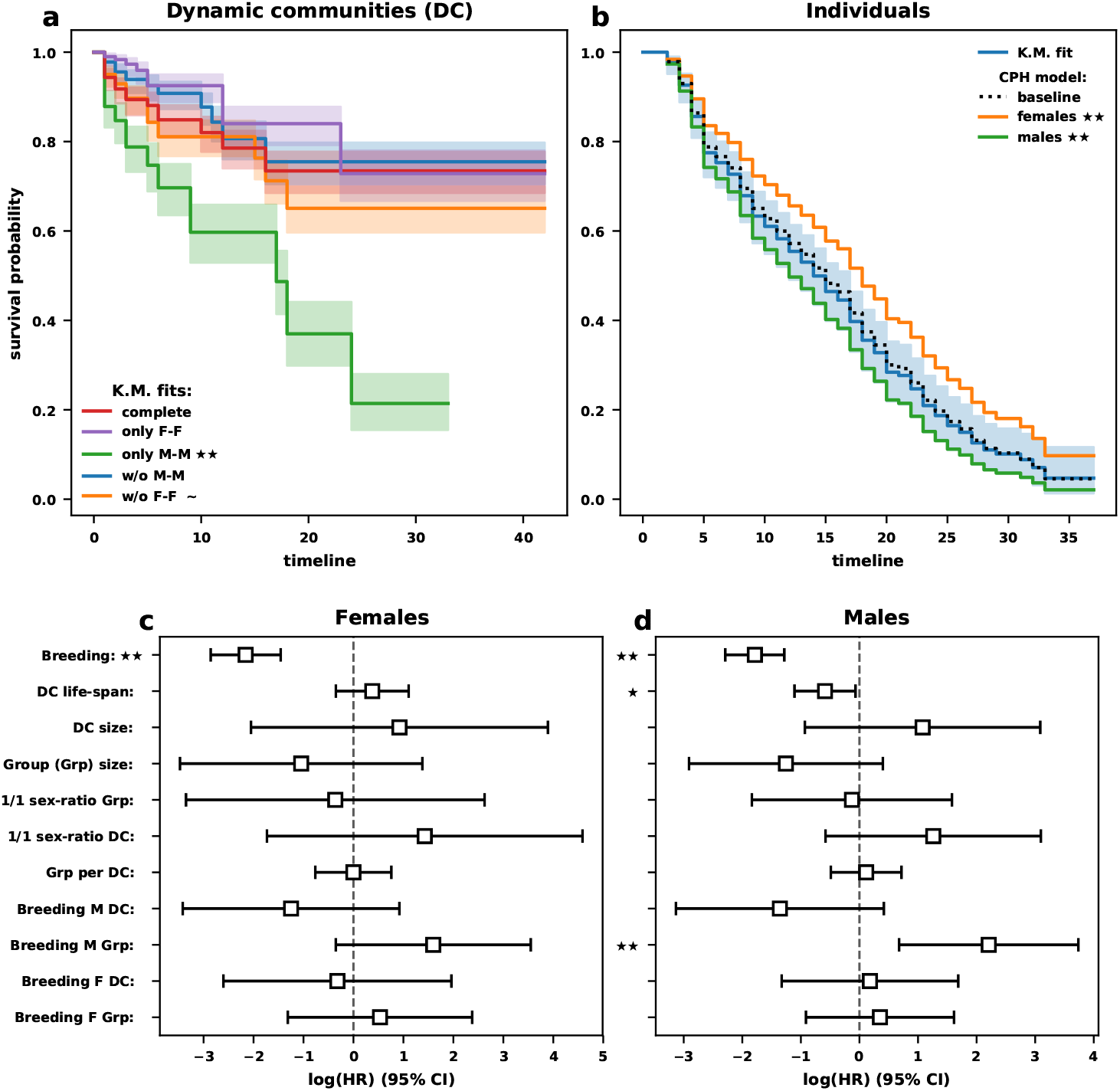
Survival analysis of dynamic communities (DCs) and its constituents. **a**, Survival probability of the observed dynamic communities using a Kaplan-Meier fit (Kaplan and Meier, 1958) with estimated 95% confidence interval. The community structure for filtered versions of the complete contact structure (red) are considered. Filtering consists of exclusively female-female (purple), exclusively male-male (green), all but female-female (orange) or all but male-male (blue) interactions. Both cases excluding female-female interactions show significantly shorter community life-spans, as compared to the complete connection network. **b**, Survival probability of RFID-tagged adults, to be understood as the probability of an individual to remain within the population and not as the probability of an individual to survive. A Kaplan-Meier fit is used to represent the survival data (blue line). Survival curves for females (orange) and males (green) highlight the effect of the covariate sex on the baseline survival (dotted line) predicted by a Cox proportional hazard (CPH) model (Cox, 1972). **c** and **d**, Effect of several covariates on a tagged individual’s presence time in the population under a time-varying CPH model (Andersen and Gill, 1982). Females (panel c) and males (panel d) were studied separately. Reported is the normalised effect of covariates (see Section 4.5) when fitting a model using all covariates combined. Next to each label a *χ*^2^ test indicates the significance in increase of goodness of fit when adding a covariate as last covariate to the model. Negative values indicate beneficial and positive ones decremental effects. Statistical evidence is reported in all panels as follows: ~: *p* ≤ 0.1; *: *p* ≤ 0.05; **: *p* ≤ 0.01

### 2.5 Relation between structural persistence and individual performance

To explore how the presence of spatially stable, long-lived dynamic communities affects individual performance we carry out a *survival analysis* to relate the presence time of individuals to individual and community characteristics. Note, that the term *survival* solely describes the type of analysis we use to study the presence time of individuals and does not refer to individual survival.

Fig. 4b depicts a non-parametric fit (Kaplan and Meier, 1958) of the raw presence data, the baseline hazard, along with the effect of sex on the hazard function under a Cox proportional hazard model (CPH, see Section 4.5 for details). Being male decreases the median by a factor of 2/3, versus being female. The CPH identifies a significant net negative effect of sex on the presence time (relative risk, *RR* = 1.66; *p* < 0.001) (see SI Section 2.2 for a definition of the relative risk).

Given that sex has a drastic effect on the presence time, we carry out the survival analysis for each sex separately. We analyse the presence time of females and males independently, each by means of a time varying CPH regression model (see SI Section 2.2). Considering reproduction, we add as covariates whether an individual reproduced at least once during the study period (*Breeding*) or not. Further covariates are the life-span of the current dynamic community (*DC life-span*), the size of the current group and dynamic community (*Grp size*, *DC size*) the deviation from an equal sex ratio within the group and the dynamic community (*1/1 sex-ratio Grp*, *1/1 sex-ratio DC*), and the number of breeding female and males in both the current group and the current community, (*Breeding F Grp*, *Breeding M Grp*, *Breeding F DC* and *Breeding M DC*). Fig. 4c and d illustrate the effects of each covariate on the logarithm of the hazard rate under a model fit including all covariates. Several of the tested covariates show a significant net effect on individual presence within the population (see caption of Fig. 4c and d for details). The only covariate relevant for both sexes, and the only one relevant for females, is *Breeding*, indicating whether an individual ever reproduced. This relation can readily be explained under the minimal assumption that longer presence increases the chance for reproduction: a low hazard rate prolongs the presence time, which increases the chance of reproduction and thus long-lived individuals are more likely to reproduce (see also Ferrari *et al.*, 2019, who observed that females had an increasing probability to breed the longer they lived).

While no other covariate gives a net signal for females, the covariates *DC life-span* and *Breeding M Grp* show a significant relation to individual presence time of males. That the presence of breeding males negatively affects the presence of other males within the same group may result from within-group competition between males. It is interesting to add, that there is no evidence for such a negative effect of the presence of breeding males within the same community. In fact, on the level of dynamic communities, breeding males tend to affect presence of other males positively, though this signal is not statistically significant.

## 3 Discussion

We analysed temporal contact data of a population of free-ranging house mice living in a barn that allows for migration within and outside the building. Although the raw data consisted simply of RFID readings from the entrance and exit of nest boxes that the mice use for resting, hiding and rearing litters, the deduced dyadic contacts allowed to describe and correlate behavioural patterns with seasons, sex and the reproductive status of tagged adults. As such, our study showcases that a systematic collection of elementary data provides non-trivial insights on the species’ social system. We find that the house mice in the study population live in multi-female and multi-male dynamic communities. Such dynamic communities are characterised by spatial fidelity, strong temporal consistency and with a life-span extending the average presence time of an individual by 2.5 times. Communities fission in a few groups in summer, which fuse again in winter, during the off-breeding time. Contrary to expectations, female-female contacts are crucial for community stability within the population, with male-male contacts strongly affected by seasonality (breeding versus off-breeding time).

Female interactions form the basis for the long-lived nature of the observed dynamic communities. Even when considered in isolation, female-female connections lead to dynamic communities that do not, in terms of life-span, show significant deviations from what we observe in the population as a whole. This observation might help to explain why such structures exist, but it needs to be considered in the light of the chosen approach: network-based group detection, and ultimately group cohesion, do not result from frequency considerations of dyadic connections alone. They also depend on how interactions are distributed within the population, which is likely to depend on both sexes. Our results reveal that females are socially strongly confined to only meet with group members in nest boxes. Hence, we find females to be less mobile than males, in terms of community association. Only 23% of the females belonged to more than one dynamic community within their life-span in the barn, in contrast to 36% of the males. In addition, the presence time of males in the study population was generally shorter than that of females. Overall, our results suggest a driving role of female-female associations for community dynamics. This reflects recent evidence of a more active role in female behaviour in the structuring of a population (Hurst, 1987; Stockley and Bro-Jørgensen, 2011; König and Lindholm, 2012; Coombes *et al.*, 2018) Our findings further support the suggested importance of bonds among females to improve their reproductive success (Weidt *et al.*, 2014; Harrison *et al.*, 2016).

The organisation of a population into social groups, often considered to be demes in which a single male monopolises breeding access to several females, has been reported repeatedly and consistently for house mice (Bronson, 1979; Crowcroft and Rowe, 1963; Reimer and Petras, 1967; Singleton and Hay, 1983; Poole and Morgan, 1976), with an identification of the dominant male via a display of aggression directed towards subordinates (Poole and Morgan, 1976; Singleton and Hay, 1983). Despite several attempts, the available contact data did not allow for a systematic identification of connection patterns that would reveal dominance, and thus allow us to detect a dominant male within social groups. That the presence of males is negatively affected by the presence of other, breeding males within the same group is the only indication for aggression, favouring the hypothesis that males improve reproductive success by evicting other males from their territory. Thus, we find no support for the idea that a single male is the core of a social group. Instead, our analysis shows that house mice live in dynamic social communities consisting of multiple males and females with both breeding and non-breeding individuals of both sexes. Even more interestingly, we observe a trend that breeding males have a positive effect on the presence time of other males, which allows for the speculation on the existence of male associations being important for reproductive success. In this regard, further exploration of the relation between social structure and genetic relatedness (Evans *et al.*, submitted) might allow to better understand the mechanisms at hand.

On the level of an individual house mouse, summer periods - with high breeding activity in the study population - lead to fewer and more specific interactions, especially if the individual is breeding. On the population level, summer is characterized by a division into more and smaller groups, as compared to winter. The transitions between winter and summer thus entail a general restructuring of direct contacts in addition to a division and recombination of groups. In light of these findings it is not necessarily an expected observation to find dynamic communities with life-spans extending over several seasons. In fact, this is only possible if the restructuring of direct contact patterns is mainly contained within dynamic communities and thus, if communities fission into sub-groups during summer that recombine during the cold winter period. These findings allow for speculations on the effect of intrasexual reproductive competition on the structuring of groups and dynamic communities: In house mice, both males and females compete with same-sex group members over reproduction (for a summary of the evidence see Carlitz *et al.*, 2019), and this is expected to affect group structure seasonally. In winter, house mice might form larger groups because of thermo-regulatory benefits (winter huddles) and a lower potential for competition over mating partners. Nevertheless, the fact that they do not abandon their community structure during off-breeding months suggests the importance of the social environment for successful reproduction. Given that house mice can potentially breed all year round, this might allow for the rather opportunistic onset of reproduction in competitive individuals, if environmental conditions permit it (occasional reproduction during winter occurs in the study population König and Lindholm, 2012; Ferrari *et al.*, 2019).

With the average life-span of a community extending roughly 2.5 times the presence of a tagged individual, the social structure within this population can be considered remarkably stable. Both sexes disperse (Runge and Lindholm, 2018), nevertheless, females mainly remain in their community they belonged to when tagged as young adults, which results in spatial stability of dynamic communities over long periods of time. This might be expected in species where females compete over areas that provide resources needed to rear their young, and where males compete over access to areas where the females are. In contrast, species where a dominant male directly monopolises sexually mature females, we rather expect that a group will disappear when the male dies, with females moving to another male/group (Clutton-Brock, 2016). Our observation of stable dynamic communities (stable both in time and space) again supports the idea of an important role of female-bonded groups in the structuring of a population.

In conclusion, we note that our observations, by means of tracking data, allowed the identification of previously unknown population structures along with evidence for their importance on individual presence within the population of house mice. We were able to demonstrate the importance of female-female associations for the stability of dynamic communities. Although males have a drastically shorter presence time in the barn than females before they leave the building or die, belonging to a stable dynamic community and interacting with other breeding males within it increases their duration of presence and thus, potentially, their reproductive success. Nonetheless, our analysis also highlights the limitations of a purely automated approach. We feel that automated tracking is most valuable if it can be combined with more traditional observation methods. In particular, direct observations should readily allow to identify directed aggression between males. And observations of male-male interactions within and between dynamic communities would allow to test the hypothesis of male cooperation within dynamic communities.

## 4 Materials and Methods

### 4.1 The house mouse population

The house mouse study allowed a longitudinal observation of a free-ranging population of individually marked house mice (König and Lindholm, 2012; König *et al.*, 2015; Geiger *et al.*, 2018; Ferrari *et al.*, 2019). The experiment was set up in 2002 in a barn of approximately 72m2 near Zürich, Switzerland, where a free-living population of *Mus musculus domesticus* resides until today. The inside of the barn is depicted in Fig. 1a. Founded with 12 individuals captured on surrounding farms, the population census reached over 600 individuals in recent years (Geiger *et al.*, 2018; Ferrari *et al.*, 2019). Food is replenished every two to three days and thus available *ad libitum*. This imitates the natural habitat in which house mice have evolved during the last several thousand years, in or near human settlements (farms, stables, storage buildings, etc.) with abundance of food. The data used in our study originates from an automated system installed in 2007 (for details seee König *et al.*, 2015). Mice are tagged with a radio-frequency identification (RFID) chip once they reach a minimum weight of 18 grams. Fourty artificial nest boxes are spread out equally in four larger, connected areas. Each nest box can be accessed through an entry tube that is equipped with two RFID antennas (for a schematic representation see Fig. 1c). Combining information from both antennas allows to record the mice’s movements in and out of boxes. With a diameter of 15 centimeters nest boxes can accommodate several individuals at once, and as many as 28 tagged mice were observed on a single occasion in the same nest box during the observation period.

Mice are free to move between the four larger areas and to enter and exit the barn. Even though possible, immigration is very rare and mainly consist of single events where offspring from a previously emigrated female returns into the barn (unpublished observation). Emigration out of the barn is common among subadults, before individuals were tagged (Runge and Lindholm, 2018).

The study needs two types of routine checks. Roughly every 10 days, there is a nest check to account for new litters (König and Lindholm, 2012; Auclair *et al.*, 2014b,a). Pups are sampled (piece of tissue taken by ear punch) in order to determine genetic maternity and paternity using 24 microsatellite markers (Auclair *et al.*, 2014a; Harrison *et al.*, 2018; Ferrari *et al.*, 2019). Approximately every two months, there is a general population check, during which all individuals that reached 18 grams are transpondered with an RFID chip, and again genetically sampled. In the present study, we use that information to characterise an individual as breeding or not, see below. The tracking system provides no information on the movements of mice before they are tagged.

An individual is considered *breeding* as soon as there is genetic evidence of it having bred once. We subtract three weeks for gestation (19-21 days) from the estimated date of birth of the progeny to define the onset of breeding. The date of birth of the progeny is estimated based on the pups’ morphological development at the day of first found (with an accuracy of ±1 day), given the frequent routine check for litters (for details see Auclair *et al.*, 2014a; Ferrari *et al.*, 2019).

We choose to study the data from January 2008 to January 2010. During this period, the antenna system functioned robustly and the population size increased from approximately 60 to a maximum of 150 individuals.

### 4.2 Deduction of pairwise contacts and the resulting contact network

The systematically collected movement data from the RFID-system was used as elementary data points to reconstruct the population-level structure and dynamics of this house mouse population. Nest boxes are too small for individuals not to register the presence of others and thus extended simultaneous presence requires the involved individuals to tolerate each other. Hence, we use the time individuals spend together in a nest box as indication for their social connection. More precisely, we consider the total time spent concurrently in a nest box as a measure of association strength between individuals. This is independent on whether, and if how many, other individuals are also present during the concurrent stay. If more than two individuals reside simultaneously in a nest box then the association strength increases for all pairs of the present individuals. As such, the resulting contact or association strength between a pair of individuals should not be seen as a direct measure for active pairwise bonding, but rather as an indication for the absence of any aggression between the two individuals. Aggregation of pairwise contact time over a fixed period (14 days in this case) then allows us to create a network representation of the contact structure of the entire population.

To account for temporal changes, we create a time-series through repeated aggregation over equally long, non-overlapping time-windows. Inactivity periods of the data collection system (indicated as grayed vertical bars in Fig. S1) prolong the affected time-window, such that each aggregation window consists of an equally long period of active data collection, and not necessarily of effective time passed. Note, that at a tagging event (approx. every two months) the data collected during 24h around the event is ignored, in order to exclude the potential disturbance it causes. The resulting time-series of contact networks, also called time-window graphs (Holme, 2015), is a dynamic representation of this population of wild house mice.

### 4.3 Detection of seasonal differences

To test for seasonal patterns, we consider a summer period (approx. July to September) and a winter period (approx. January to March). More precisely, we consider five aggregation windows, starting once in January and once in July in each of the two years, 2008 and 2009. The summer and winter periods are highlighted in red and blue, respectively, in Fig. 1d. For measures taken at the population level, like sex-ratio or contact density, we consider the 10 aggregation windows per season as independent replicates. We use a two-sided Kolmogorov-Smirnov test to assess the significance level of seasonal difference in sex-ratio.

Measures taken on the individual level, like contact specificity, are considered per individual, and thus are averaged over all aggregation periods in which the individual of interest is present. These measures are reported per individual per season.

To assess how partner-specific an individual spends its time in contacts, both in summer and in winter, we normalise the individual’s total pairwise contact time and generate a cumulative contact time distribution, ranked from the most to the least important contact during an aggregation window. We average the cumulative distributions over the summer and over the winter periods for each individual to assure that each individual contributes equally. We would argue that the steeper the assent, i.e. the bigger the fraction of contact time attributed to the most important contacts, the more specific to the choice of contact partners an individual is. We classify the cumulative distributions of all individuals according to sex of the target individual (male/female) and the season (summer/winter). Combining all distributions within each class results in 4 datasets: female-winter, female-summer, male-winter and male-summer. Each dataset holds, for each contact partner, ranked in position from most to least important, a distribution of contact time fractions attributed to this and more important contact partners. For each position, a two-sample t-test is carried out to determine whether the association of contact time differs between summer and winter.

### 4.4 Assessment of population structure and dynamics

We use partitioning of the contact network to describe its structure. Partitioning is a separation into groups of individuals with stronger within-group contact density. For every slice, the partitioning process consists of two steps: first, the contact network is decomposed in a set of disconnected sub-networks, i.e. each set being an ensemble of individuals with connections exclusively inside the set; second, within each sub-network a standard network partitioning method, the heuristic algorithm by Rosvall and Bergstrom (2008), is applied to further partition the sub-networks. Combining both steps leads to a partitioning of the population into denser connected groups.

The partitioning algorithm applied in the second step tries to partition a network into groups of nodes that predominantly connect between each other. The principle underlying this algorithm can best be understood by imagining a process that moves from node to node following connections: The algorithm favours group configurations for which this process crosses group borders less often. Ultimately, the heuristic procedure suggests a group structure where between group transitions are as rare as possible, leading to a group structure that maximises within group movements of the process and hence within group connections. If the contact structure does not present regions with more densely connected nodes, then the algorithm might put all nodes in the same group (Lancichinetti and Fortunato, 2009). Thus, neither the first, nor the second step in the procedure applied here enforces a partitioning.

By partitioning the contact networks in each slice, we construct a time-series of decompositions of the contact structure within this population. The time-series of decompositions is, in fact, a representation of the dynamics of the structural organisation within the population. A description of the structural dynamics can thus be achieved through a comparison of the group structure between different points of the time-series, a task that qualifies as dynamic community detection (Holme, 2015; Dakiche *et al.*, 2019). We utilise the evolutionary clustering method by Liechti and Bonhoeffer (2019) to detect dynamic communities. This method detects dynamic communities by linking groups between different time points. If two groups from different time points consist of the majority of members of each other, then the algorithm considers them as temporally different representations of the same community. Based on the ensemble of these representational relations, the algorithm then constructs the time course of dynamic communities. The advantage of this procedure is that it can detect time persistent structures that are not necessarily continuously expressed. In particular, any sort of temporary decomposition of a community might not lead to its dissolution: If two groups from distant time points are representations of the same community, and the considered members do only regroup among each other in the group structures in-between, then the algorithm can still pick up the over-arching community structure. How temporally distant two groups can maximally bridge such decomposition-gaps is determined *a priori* through a parametrisation of the method. This approach allows the method to detect dynamic communities with internal fission-fusion dynamics. To the best of our knowledge, there is currently no other dynamic community detection method with a comparable feature available.

The sole parameter required by the method, the *history parameter*, determines the maximal duration over which groups can be considered as temporally different representations of the same dynamic community. While one might freely choose its value, we use a 12-step history, a value that maximises the *total consistency* within the dynamic community structure in our study population (see Liechti and Bonhoeffer, 2019, for details).

### 4.5 Survival analysis of dynamic communities and individuals

All survival analysis presented in this study were carried out using the implementations in the software package *lifelines* by Davidson-Pilon *et al.* (2019).

To study the contribution of sex-specific contacts to the longevity of the observed dynamic communities, we construct series of contact networks once ignoring and once exclusively considering same sex contacts. Based on these networks, we determine dynamic communities and carry out non-parametric fits (Kaplan and Meier, 1958) for the resulting survival data. Caution needs to be taken as the sample size of dynamic communities in the survival analysis is not fixed, but rather depends on the actual time of existence (life-span) of each sample. This stems from the combination of observing over a fixed duration and the fact that the population is categorised into dynamic communities at each time point. Consider the case where all individuals reside in a single long-lived community. If this community persisted throughout the observation period, we would be left with a single observation. If it existed only for half a year and were replaced by a sequence of dynamic communities with a life-span of one month each, for the rest of the year, our samples would mainly consist of short-lived dynamic communities. This would occur even though, throughout a year, short-lived dynamic communities would be observed equally often as long-lived ones. To account for this bias, we consider the presence of a community at each point in the time-series as an observation. Thus, for a community spanning over the entire observation period (42 aggregation windows), we would report 42 right censored observations rather than just a single one. To assess the statistical significance of the eventual differences observed we perform pairwise logrank tests (Mantel, 1966).

On the individual level, we measure an individual’s presence time in numbers of slices (aggregation windows) between the points where an individual was first and last observed. Only individuals are considered that appear later than the first time point in the time-series of contact networks. All individuals present at the last time point are right censored. To assess how presence time relates to sex, we fit a Cox proportional hazard model (Cox, 1972, or refer to SI Section 2 for further information) with sex as only covariate. We consider Schoenfeld residuals (Schoenfeld, 1982) to assert no violation of the proportional hazard assumption (results not shown).

Studying the entire population, as well as the stratification by sex, bares the risk to convolve effects of covariates that are sex-specific. As a consequence, we analyse the within-population presence for females and males separately. Since individual properties, like the breeding status or number of other breeding males within the group, change over the life history of an individual, we also switch to a time varying version of the Cox proportional hazard regression model (see Andersen and Gill, 1982, or refer to SI Section 2.3 for further details). We perform a series of model fits including different combinations of the above covariates. To assess the gain in goodness of fit when adding a covariate, the change in log-likelihood is evaluated using a *χ*^2^-test when adding this covariate as last covariate to the model. The significance of these tests is reported next to the name of each covariate in Fig. 4c and d. Each covariate is normed, either by its average or by an expected value. This normalisation does not change the effect of a covariate, it is done only to allow a combined display of the covariates. Without this harmonisation of the magnitudes it would be difficult to jointly report covariates. The effects of a covariate on the logarithm of the hazard rate, as illustrated in Fig. 4c and d, can be exponentiated to yield the relative risk factor of each covariate. Hence, negative values indicate beneficial and positive ones decremental effects. We speak of a net effect if the 95% confidence interval is homogeneous in the direction of the effect.

## Supporting information

Animated average poistions with contacts (barn_anim.mp4).

## 5 Acknowledgements

We like to thank Hopi Hoekstra and the entire Hoekstra lab, especially Nicholas Jourjine, for the many valuable feedbacks they provided on this manuscript. We would also like to thank the numerous helpers who contributed to the maintenance of the study site and to data collection, here especially Anna Lindholm and Jari Garbely. Further thanks go to Barbara Mejìa for the detailed feedback on structure and style of the manuscript.

Financial support was provided through the SNF grant 310030B 176401 to SB, the SNF grant 31003A 176114 to BK and by the University of Zurich.

## 6 Ethical statement

Methods described and data sampling of the study population were collected according to the Swiss Animal Welfare Ordinance (TSchV), with permit approved by the Veterinary Office Zurich, Switzerland (Kantonales Veterinäramt Zürich, no 215/2006).

## Supplementary Materials

### SI 1 Supplementary figures

**Fig. S1.**
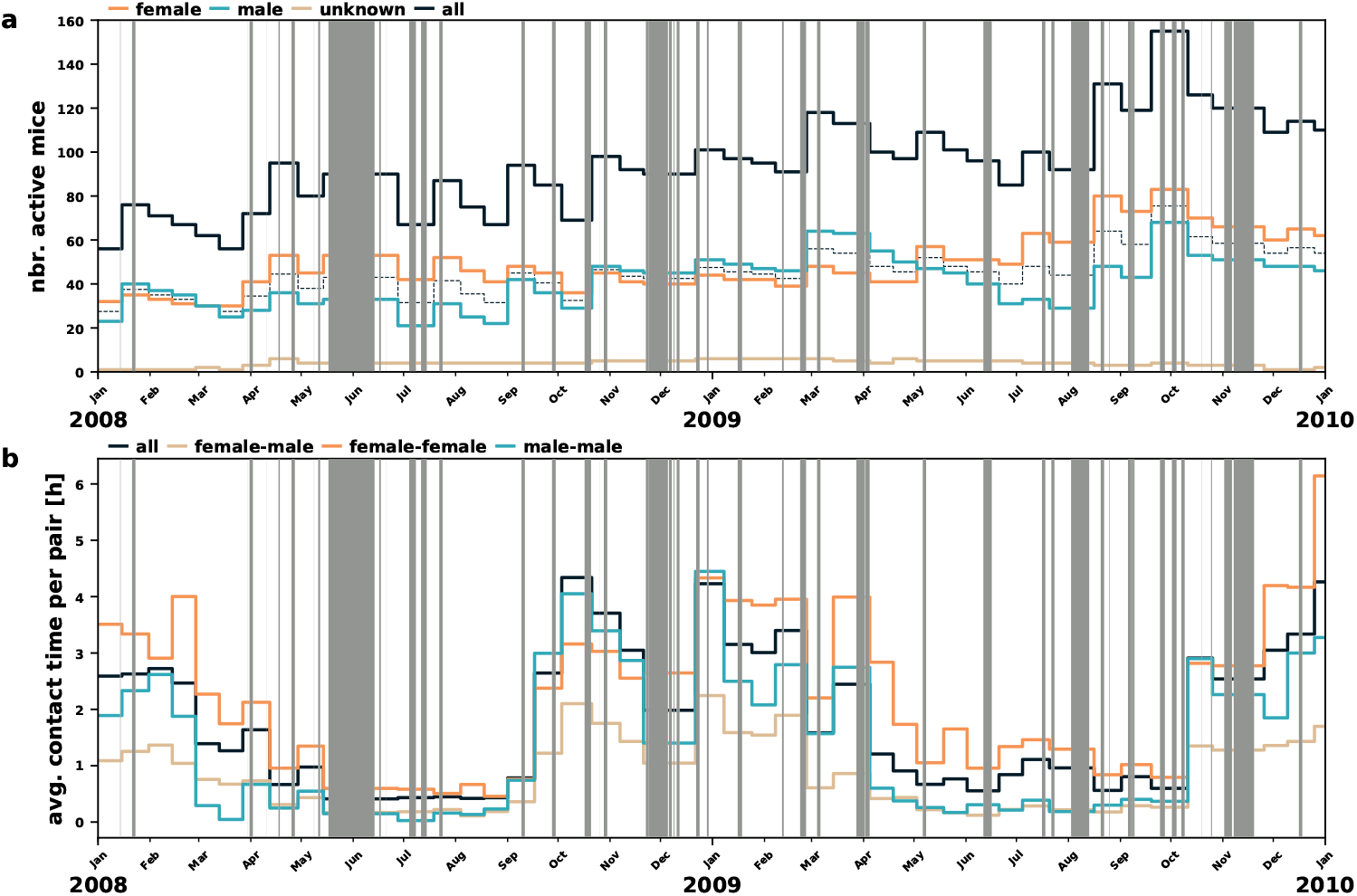
Overview of system down times during the data collection period. Grayed vertical bars indicate periods with no data collection. These periods were removed during the analysis and re-introduced afterwards. **a**, Time-series of the number of house mice detected by the RFID-readers during consecutive observation periods of 2 weeks. The data shown here is equivalent to Fig. 1d. **b**, Average contact duration between individuals from different categories. The data shown here is equivalent to Fig. 2b.

**Fig. S2.**
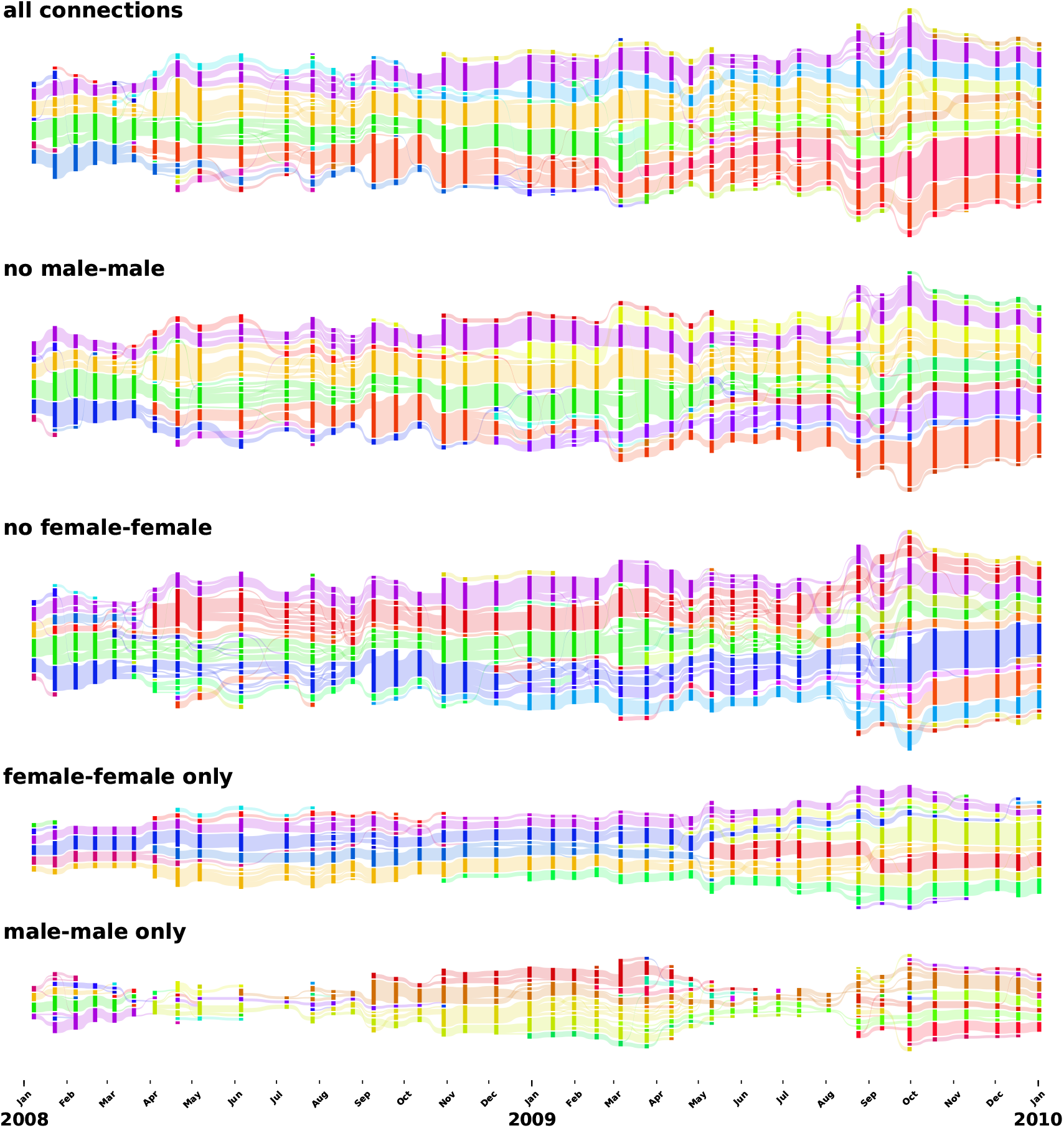
Illustration of the dynamic community structure in filtered representations of the contact data. The colouring of dynamic communities is drawn from the same sequence of colours for each illustration. Consequentially, the colouring is not consistent between the alluvial diagrams, as not all representations produce the same ordering of dynamic communities. Note, however, the similarity in colouring between the top most panels reflects the existence of only marginal changes in the dynamic community structure when ignoring male-male connections. The alluvial diagrams were created in python, using the *pyAlluv* package (Liechti, 2020).

**Fig. S3.**
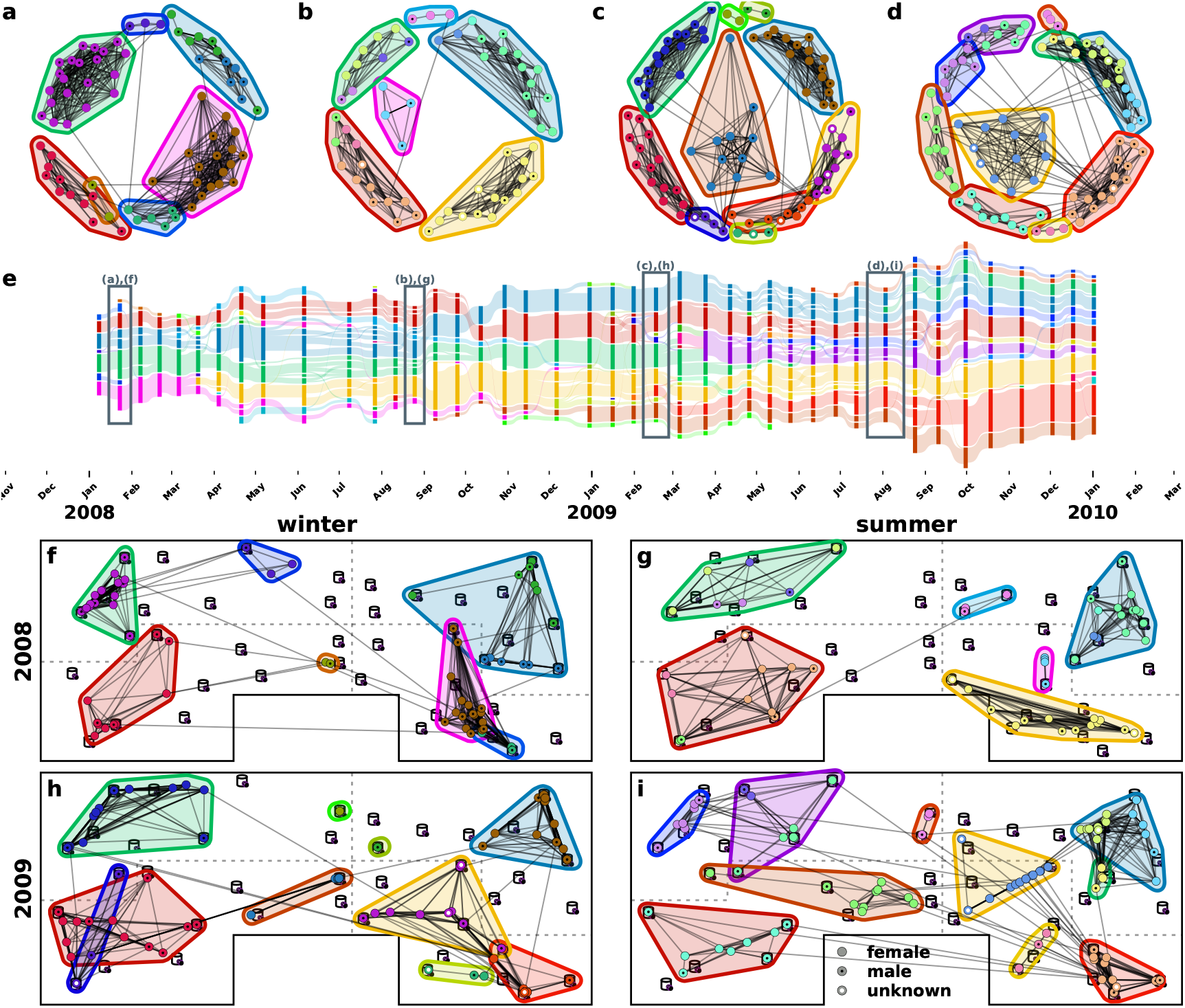
Temporal and spatial representations of the contact structure. **a** to **d**, Representations of the contact network at randomly chosen time points during the summer and the winter periods. The layout is deduced using the force-directed spring model by Fruchterman and Reingold (1991); Hagberg *et al.* (2008). Node colours indicate the group structure. The contours represent the classification of these groups into dynamic communities. The colouring of the contours is consistent throughout the figure with the life-cycle of each dynamic community depicted in panel (e). **e**, Alluvial representation of the dynamic communities. The social and spatial illustrations depicted in the other panels are marked with a gray box. **f** to **i**, Illustration of contact structures embedded within the barn. Individual house mice, i.e. dots, are placed at their average position in the barn, i.e. the barycentre of the nest boxes they frequented during the aggregation window weighted by the total time spent in each box. Node colours match the corresponding network representation in panels (a) to (d).

**Fig. S4.**
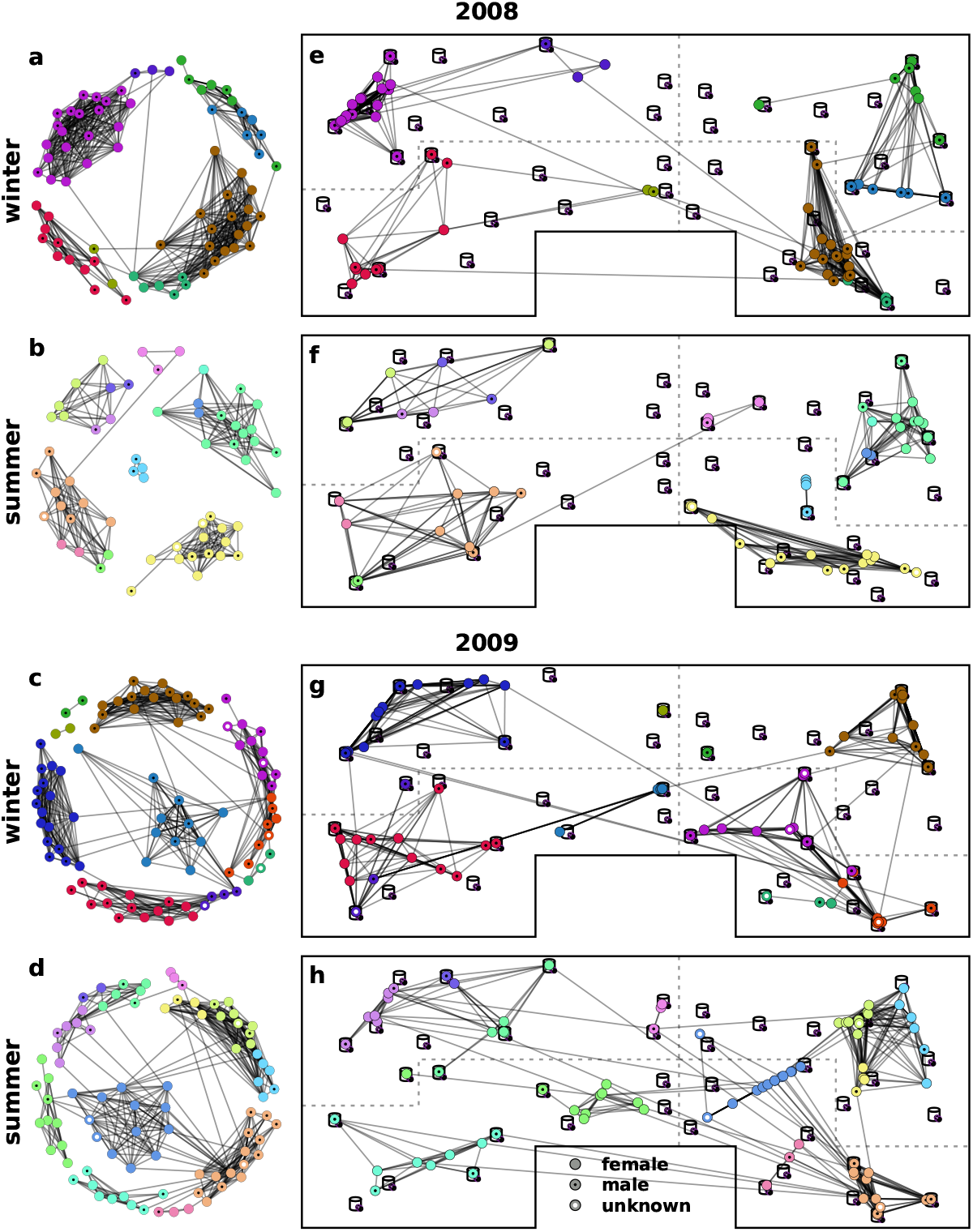
Temporal and spatial representations of the contact structure. Colours indicate different groups found by a partitioning analysis (see Section 4.4 for details). **a** to **d**, Representations of the contact network at a randomly chosen time points during the summer and the winter periods. The layout is deduced using the force-directed spring model by Fruchterman and Reingold (1991); Hagberg *et al.* (2008). Node colours indicate group structure. **e** to **f**, Illustration of contact structures embedded within the barn. Mice are placed on the weighted barycentre of the nestboxes they frequented during the aggregation window. Node colours match with the corresponding network representation in panels (a) to (d).

### SI 2 Cox proportional hazard

#### SI 2.1 Hazard function

Survival analysis is the study of the temporal persistence of certain objects or states and the putative relation between this persistence and other covariates. The basic approach in survival analysis is to assess and study the hazard function, a function that describes the change in probability to exist, given a certain age. For a set of objects we can define the hazard curve as follows:

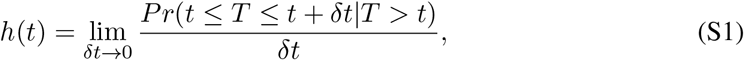

where *T* is the time of persistence of a randomly chosen object.

#### SI 2.2 Cox proportional hazard regression model

The Cox proportional hazard regression model (Cox, 1972) allows to study the effect of a set of covariates on the hazard curve. It does so assuming that for a single object the logarithm of the hazard curve is a linear combination of the set *x* of *n* covariates and a baseline curve that is independent on the covariates.

In mathematical terms:

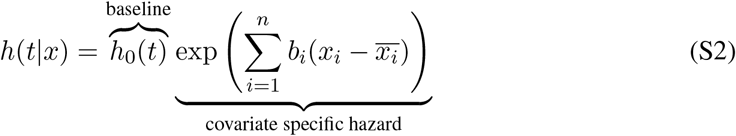

The terms *b*_*i*_ and 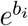 define the effect, as reported in Fig. 4, and the *relative risk* (RR) of covariate *x*_*i*_.

The regression is based on a likelihood optimisation and independent on the ordering of covariates. The regression procedure will not be detailed further (see Cox, 1972, for detailed explanations).

#### SI 2.3 Time varying Cox proportional hazard regression model

In the proportional hazard model defined before, only the baseline hazard curve changes throughout the lifetime of an object. This is a strong assumption and does not allow to include covariates that change throughout the lifetime. The time varying Cox proportional hazard model, presented in Andersen and Gill (1982), allows to overcome this limitation through the introduction of a time varying covariate, *x*_*i*_ → *x*_*i*_(*t*), in Equation S2.

